# Molecular basis for cellular compartmentalization by an ancient membrane fission mechanism

**DOI:** 10.1101/2025.11.28.690958

**Authors:** Javier Espadas, Diorge P. Souza, Markku Hakala, Juan Manuel García-Arcos, Joshua Tran, Ashutosh Kumar, Carlos Marcuello, Andrea Merino, Adai Colom, Stefano Vanni, Marko Kaksonen, Buzz Baum, Christopher P. Toret, Aurélien Roux

## Abstract

The emergence of cell compartmentalization depends on membrane fission to create the endomembrane compartments. In eukaryotes, membrane fission is commonly executed by ESCRT-III, a protein complex conserved in all domains of life. However, whether membrane fission was an ancestral ESCRT-III activity predating eukaryogenesis remains unknown. Here we show that ESCRT-IIIA from Asgard Heimdallarchaeota, the closest archaeal relatives of eukaryotes, performs membrane fission through an N-terminal amphipathic helix, which we term Hofund. In eukaryotes, Hofund is fragmented across ESCRT-IIIA paralogs, and disrupting these regions causes severe fission defect in yeast. Remarkably, Heimdallarchaeota Hofund restores fission when fused to defective eukaryotic paralogs. These findings suggest that ESCRT-III-mediated fission arose before eukaryogenesis and later diversified to support the regulatory complexity of eukaryotic compartmentalization.

## Main Text

Compartmentalization is a hallmark of eukaryotic life, enabled by dynamic lipid membranes that segregate biochemical reactions while permitting selective exchange of biomolecules. Membrane fission is fundamental to this organization. In eukaryotes, this process is commonly executed by ESCRT-III, a ubiquitous membrane fission complex operating across a majority of cellular membranes (*1*, *2*). ESCRT-III remodels membranes during cytokinetic abscission (*3*, *4*), the formation of intraluminal vesicles (ILVs) for cargo degradation (*5*), lysosomal (*6*, *7*) and plasma membrane repair (*8*, *9*), and nuclear envelope reformation (*10*, *11*). ESCRT-III filaments undergo stepwise structural changes mediated by subunit turnover upon ATP hydrolysis by the AAA+ ATPase Vps4, which increases filament and membrane curvature, ultimately thinning and cleaving narrow membrane necks (*12*, *13*). In the hyperthermophilic Crenarchaea, cell biology experiments support an essential role for ESCRT-III paralogs in membrane remodeling related functions such as cell division and release of extracellular vesicles (*14–19*). However, the mechanistic contribution of ESCRT-III to these processes, particularly whether and how these proteins sever membranes, remains to be defined. Hence, despite an ancient origin in the last universal common ancestor (LUCA) (*20*) and the widespread distribution of ESCRT-III proteins in bacteria and archaea, the ancestral biophysical activity of the ESCRT-III complex remains unresolved.

Asgard archaea provide a simplified system to probe the fission capacity of non-eukaryotic ESCRT-III proteins, as they encode ESCRT-III members and the AAA+ ATPase Vps4 (*21–23*). In Asgard Heimdallarchaeota (Heimdall), the closest known relatives of eukaryotes (*21*, *24*), the ESCRT-III machinery consists of only two subunits: ESCRT-IIIB, a “before-type” (B-type) ortholog of the eukaryotic paralogs Chm7, Vps20, Vps60, and Snf7, that initiates membrane remodeling, and ESCRT-IIIA, an “after-type” (A-type) ortholog of the paralogs Vps2, Vps24, Did2, and Ist1, that promotes membrane curvature (*25*). Heimdall also encodes a single Vps4 paralog, *ah*Vps4. Despite its simplicity, the Heimdall system retains hallmark features of the eukaryotic ESCRT-III machinery, including sequential polymerization and curvature generation (*25*). Yet whether this archaeal system is fission-competent, or instead operates solely as a membrane curvature generator, remains unknown. ESCRT-IIIA and *ah*Vps4, whose eukaryotic counterparts execute membrane fission, therefore represent ideal candidates to test whether fission by ESCRT-III predates the emergence of eukaryotic endomembranes.

Here we show that Heimdall ESCRT-III promotes fission, and therefore it provides a uniquely tractable system to dissect the fission mechanism at the molecular level. In eukaryotes, ESCRT-III-dependent fission has been documented across many endomembrane pathways, yet the underlying molecular mechanism remains unresolved. A major obstacle is the complexity of eukaryotic ESCRT-III assemblies, which comprise numerous paralogs (eight in yeast, twelve in mammals, and more than twenty in some plants (*26*)). Conflicting observations from different systems have further blurred the definition of a minimal fission machinery. Several combinations of A-type paralogs have been implicated in fission (*27–29*), but their individual contributions remain debated. Adding to this complexity, the curvature context appears inverted across experimental systems: in cells, ESCRT-III acts at the interior of negatively curved membrane necks, whereas *in vitro* reconstitutions using bacterial (*20*, *30*, *31*), archaeal (*25*, *32*), and eukaryotic ESCRT-III proteins (*33–36*) consistently show preferential assembly on positively curved membranes. Reconciling these observations has remained a central challenge in the field.

### ESCRT-IIIA promotes membrane fission upon ATP hydrolysis by *ah*Vps4

To determine whether Heimdall ESCRT-III proteins can mediate membrane fission, we reconstituted their activity *in vitro* using purified components from the Heimdallarchaeon AB_125. By analogy to eukaryotes, where A-type subunits execute fission, we hypothesized that Heimdall ESCRT-IIIA would carry the fission activity (Fig. 1A). To test this, we first established an assay to generate lipid nanotubes (NTs) suspended between silica beads, enabling real-time visualization of membrane dynamics and ESCRT-IIIA recruitment (Fig. 1B, top/mid panels). Like eukaryotic A-type subunits, ESCRT-IIIA bound efficiently to positively curved NTs but failed to associate with negatively curved membranes (fig. S1A,B). A curvature-sorting analysis showed strong enrichment onto NTs above 0.023 nm⁻¹ (radius ≈ 31 nm), comparable to the behavior of human ESCRT-III (*36*) (fig. S1A,C). Upon binding, ESCRT-IIIA formed dynamic puncta (fig. S1D,E) that matured into stable assemblies, that locally constricted NTs at pH 6, reducing radii from ∼35 nm to ∼10 nm (Fig. 1C-F), demonstrating that ESCRT-IIIA can both sense and generate positive membrane curvature.

**Figure 1.**
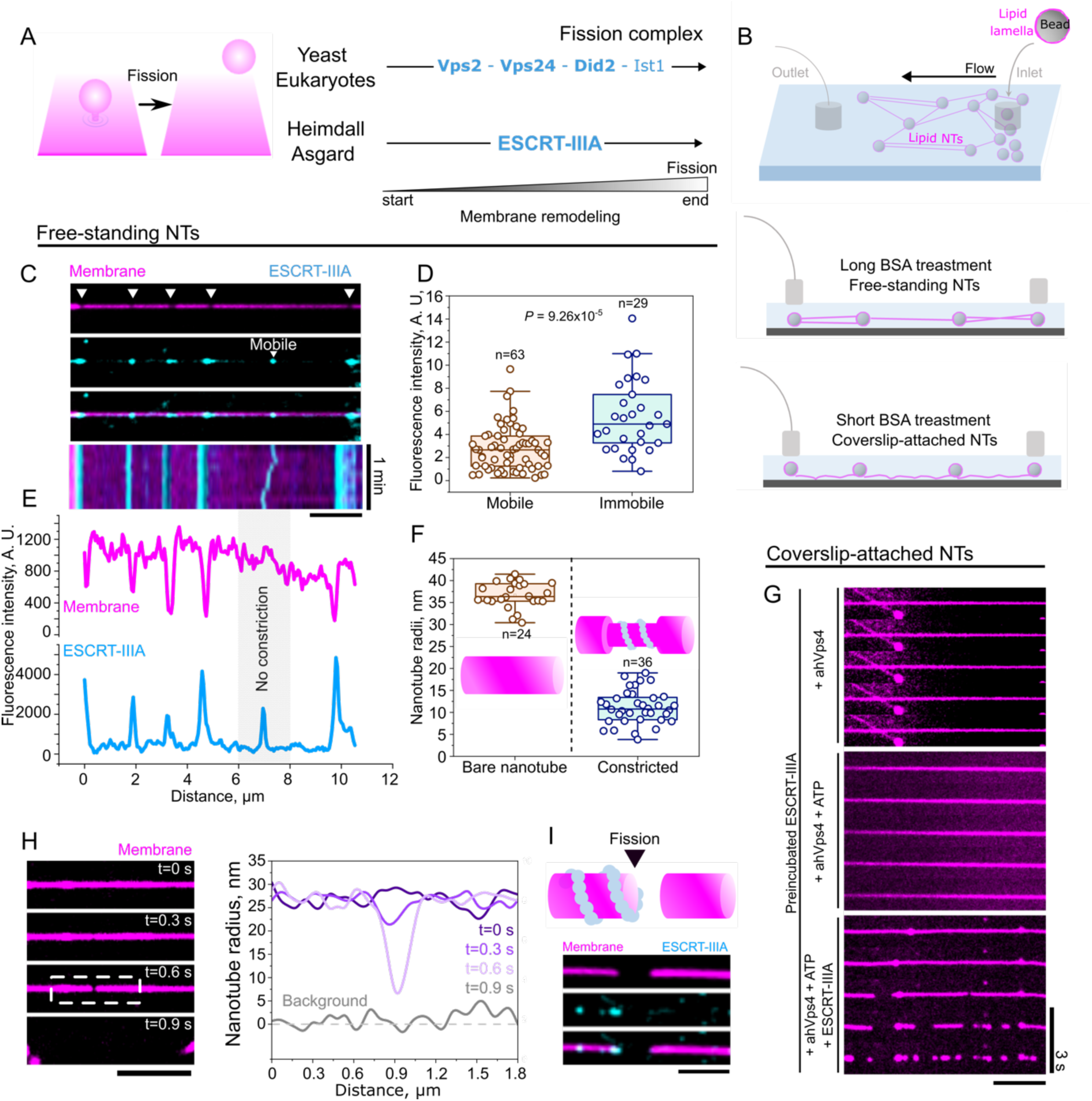
Heimdall ESCRT-IIIA drives membrane constriction and fission *in vitro*. (**A**) Conceptual model illustrating how Heimdall ESCRT-IIIA recapitulates the function of eukaryotic A-type subunits to remodel and sever membranes. (**B**) Schematic of the microfluidic setup used to reconstitute membrane fission on lipid nanotubes (NTs). Long or short coverslip-passivation times yield free-standing (mid panel) or surface-adhered NTs (lower panel), allowing real-time visualization of protein self-assembly and of fission dynamics, respectively. (**C**) Fluorescence images and kymograph showing immobile ESCRT-IIIA assemblies (cyan) constricting membrane NTs (magenta) at pH 6 in the absence of *ah*Vps4 and ATP. Scale bar is 2 μm. (**D**) Quantification of fluorescence intensity distributions of mobile versus immobile ESCRT-IIIA assemblies. Two-tailed Student’s t-test with Welch’s correction for unequal variances. (**E**) Representative longitudinal fluorescence line-scans of ESCRT-IIIA (cyan) and membrane (magenta) along the NT axis. (**F**) NT radius measurements comparing bare membranes to ESCRT-IIIA–constricted regions, which confirms active tube narrowing. (**G**) Reconstitution of ATP-dependent membrane fission with coverslip-adhered NTs. Preassembled ESCRT-IIIA on NTs (1 µM) was incubated with (i) *ah*Vps4 (5 µM; top panel), (ii) *ah*Vps4/ATP (5 µM/10 mM; mid panel), or (iii) ESCRT-IIIA/*ah*Vps4/ATP (1 µM/5 µM/10 mM; bottom panel). Scale bar is 2 μm. (**H**) Free-standing NTs revealed sequential constriction events that culminate in membrane fission. The plot profile, which shows the membrane NT radius on its longitudinal axis during ESCRT-IIIA-mediated fission. Scale bar is 2 μm. (**I**) Schematic (top) and corresponding micrograph (bottom), which shows membrane scission occurring at the edge of the ESCRT-IIIA coat, where curvature and elastic stress are maximal. Membrane, magenta; ESCRT-IIIA, cyan. Scale bar is 1 μm.

Membrane necks narrowed to radii of ∼3 nm enter a regime where the energy barrier to fission becomes comparable to thermal fluctuations, making spontaneous fission possible (*37*). However, these radii are far below the membrane curvature induced by ESCRT-IIIA alone (Fig. 1E,F). This suggested the requirement for additional constriction through an external energy input. In eukaryotes, ATP hydrolysis by Vps4 supplies the energy required for ESCRT-III subunit exchange and filament remodeling (*12*, *28*). Hence, we investigated whether ATP hydrolysis would facilitate membrane fission by ESCRT-IIIA. To detect fission events, we used lipid NTs partially adhered to the coverslip (Fig. 1B, top and bottom panel), preventing NT retraction after fission. No fission occurred when *ah*Vps4 or *ah*Vps4/ATP was added to NTs pre-coated with ESCRT-IIIA, indicating that ATPase-driven disassembly is insufficient. Strikingly, fission occurred only when preassembled ESCRT-IIIA was supplied with *ah*Vps4, ATP, and free ESCRT-IIIA in solution (Fig. 1G), which indicates that ATP-dependent subunit turnover, not filament disassembly, triggers membrane fission by ESCRT-IIIA. On surface-adhered NTs, fission proceeded without detectable local increase of membrane curvature. However, in suspended NTs where we can monitor membrane shape changes, we observed a sharp local curvature increase at the fission site (Fig. 1H), revealing a two-step mechanism: ESCRT-IIIA first constricts before triggering fission at the polymer edge (Fig. 1I), where membrane elastic energy is maximized, analogous to dynamin-mediated fission (*38*).

Together, these experiments provide the first direct evidence that an Asgard ESCRT-III protein is inherently fission-competent. Heimdall ESCRT-IIIA executes membrane fission using its own polymer dynamics and ATPase-mediated subunit turnover. This establishes the minimal biochemical requirements for ESCRT-III-driven fission and demonstrates that membrane fission is an intrinsic, ancestral activity of ESCRT-III.

### The N-terminal amphipathic helix “Hofund” drives ESCRT-IIIA-mediated fission

Lipids with negative spontaneous curvature promote hemifission intermediates and facilitate the generation of packing defects by remodeling proteins that destabilize the bilayer (*39–42*). Membrane composition strongly modulated ESCRT-IIIA-mediated fission efficiency, and fission was markedly enhanced on membranes containing 40 mol% of the negatively curved lipid 1,2-dioleoyl-sn-glycero-3-phosphoethanolamine (DOPE) (fig. S2A,B). This supported that ESCRT-IIIA-mediated fission proceeds through generation of lipid packing defects. Given that electrostatic and hydrophobic interactions at the N-termini of ESCRT-III proteins are known to mediate membrane recruitment (*43*), we considered whether ESCRT-IIIA membrane insertion would create lipid packing defects. AlphaFold3 predictions (*44*) revealed that Heimdall ESCRT-IIIA adopts the canonical α-helix ESCRT-III fold (Fig. 2A), with an N-terminal α0 helix (Fig. 2B) carrying three aromatic residues (F2, F5, and W8) aligned on one face of the helix (Fig. 2C). These residues are strongly conserved across Heimdall sequences (Fig. 2D), and their high hydrophobicity and hydrophobic moment (0.543 and 0.458) suggested a direct role in membrane insertion. This architecture parallels ultrastructural studies of human Did2/Chmp1b, the late-acting ESCRT-III subunit proposed to execute fission (*12*), where analogous phenylalanines penetrate the bilayer core (*45*), raising the possibility that F2 and F5 play an equivalent role in inserting ESCRT-IIIA into the lipid bilayer.

**Figure 2.**
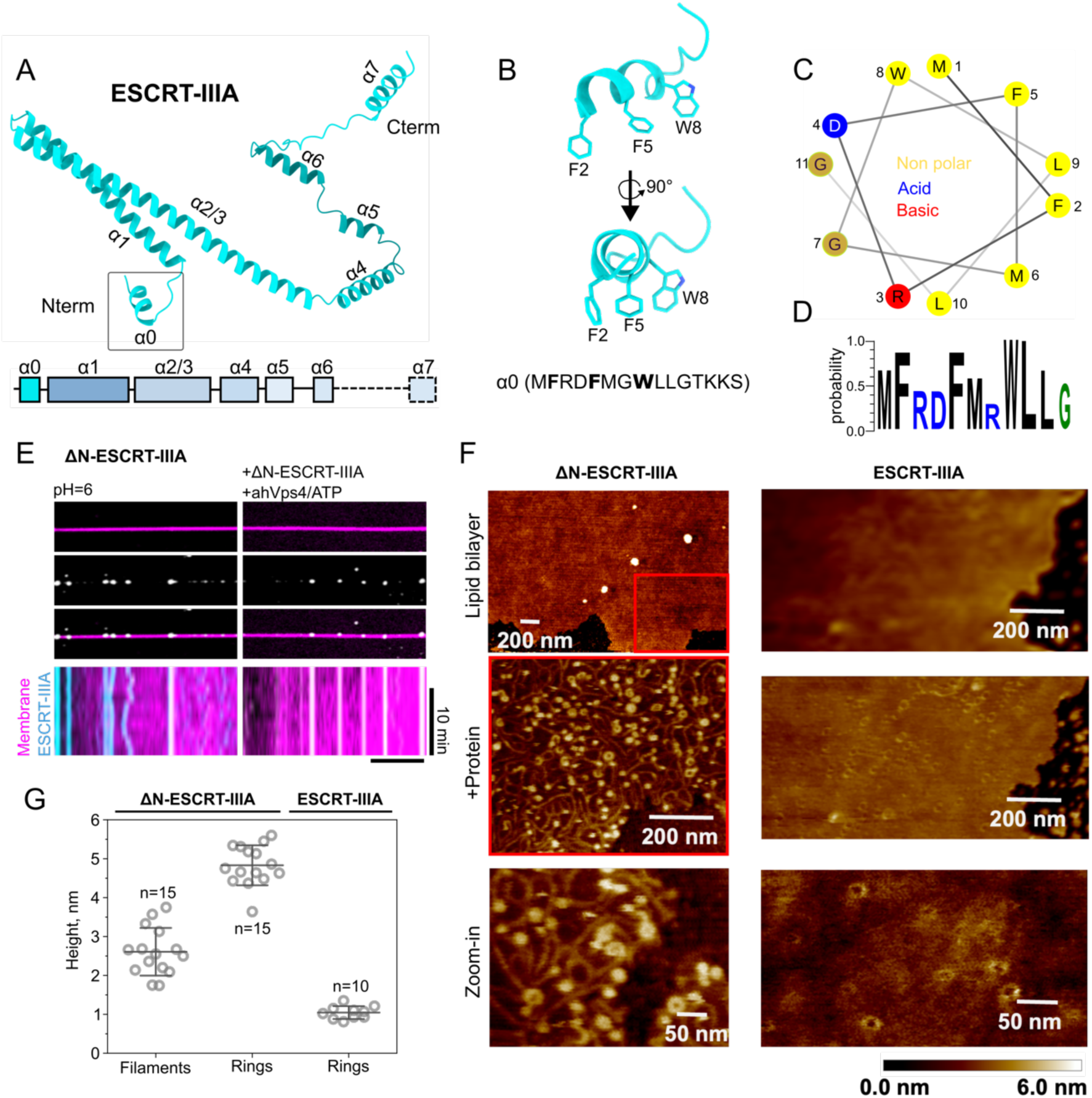
The N-terminus of ESCRT-IIIA provides the functionality for membrane remodeling. (**A**) AlphaFold3 structural prediction of Heimdall ESCRT-IIIA showing the canonical ESCRT-III fold of seven α-helices (top) and schematic representation of the primary sequence (bottom). (**B**) Close-up view of the predicted N-terminal α0 helix (first 15 amino acids). (**C**) Helical wheel diagram of the first 11 residues showing the asymmetric distribution of hydrophobic residues on one face of the helix. (**D**) Sequence logo illustrating conservation of the first 11 residues of ESCRT-IIIA across A-type subunits from Heimdall Asgard archaea. (**E**) Membrane remodeling assay using truncated ESCRT-IIIA lacking α0 (schematic representation of the primary sequence of ΔN-ESCRT-IIIA is shown at the top). Left: 1 µM ΔN–ESCRT-IIIA (cyan) incubated with NTs (magenta) at pH 6. Right: ΔN–ESCRT-IIIA preincubated with NTs followed by addition of *ah*Vps4 (5 µM) and ATP (10 mM). In both conditions, no detectable constriction or fission was observed. Scale bar, 2 µm. (**F**) AFM micrographs of *E. coli* extract lipid bilayers and of ΔN–ESCRT-IIIA or ESCRT-IIIA (2 µM) polymerized over the membrane. (**G**) Height quantification of ΔN–ESCRT-IIIA filaments and rings, and of ESCRT-IIIA over the lipid membranes from AFM height profiles.

To test whether α0 is required for membrane recruitment, we deleted the helix (ΔN-ESCRT-IIIA) and examined its recruitment and functionality on lipid NTs. ΔN-ESCRT-IIIA still localized to NTs, but failed to constrict or promote fission under conditions where the full-length protein remained active (Fig. 2E). Thus, the N-terminal α0 helix, hereafter Hofund after Heimdall’s sword in Norse mythology, is the essential mechanical trigger for ESCRT-IIIA-mediated membrane fission.

Atomic force microscopy (AFM) on negatively charged *E. coli* lipid bilayers showed ΔN-ESCRT-IIIA is capable of binding membranes and self-assemble as filaments and rings, with ring heights consistent with two stacked filament turns. In contrast, full-length ESCRT-IIIA formed exclusively low-height (∼1 nm) rings on the membrane surface (Fig. 2F,G), indicating that the α0 helix drives deep membrane insertion. Circular dichroism of the isolated N-terminal 15 aa showed that Hofund folds into an amphipathic helix upon membrane binding. The F2A/F5A mutant exhibited poor folding and substantially reduced hydrophobicity and hydrophobic moment (0.346 and 0.298) (Fig. 3A). All-atom molecular dynamics (MD) simulations revealed that Hofund inserts deeply into the bilayer, positioning F2, F5, M6, L9, and L10 ∼5 Å below the phosphate plane and embedding most of the helix within the headgroup-acyl chain interface (Fig. 3B). This insertion was entirely lost in F2A/F5A mutants, confirming that aromatic residues are indispensable for membrane penetration.

**Figure 3.**
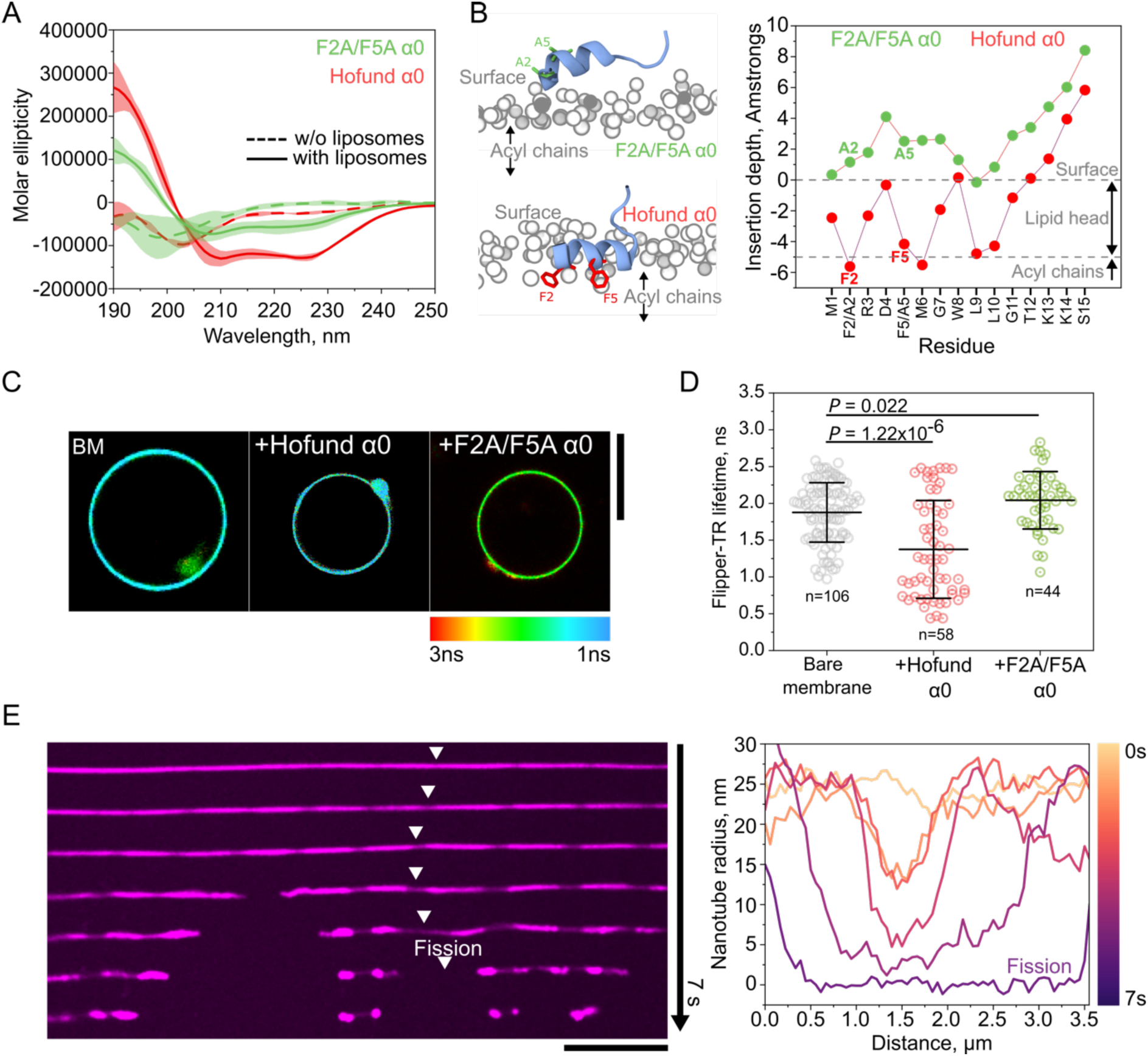
Hofund is the minimal fission machinery in ESCRT-IIIA. (**A**) Circular dichroism spectra of the Hofund α0 peptide (red) and its F2A/F5A mutant (green). Solid lines show spectra in the presence of liposomes; dashed lines correspond to buffer-only controls. The data reveal membrane-induced folding of Hofund α0 into a helix, which is strongly reduced in the F2A/F5A mutant. (**B**) All-atom molecular dynamics simulations showing membrane insertion of Hofund α0 (bottom) and its absence in the F2A/F5A mutant (top). Phospholipid headgroups are shown as gray circles. The plot (right) quantifies the insertion depth of individual residues, highlighting loss of membrane penetration in the F2A/F5A mutant. The dashed gray line indicates the membrane surface boundary. (**C**) Fluorescence lifetime imaging microscopy (FLIM) of the membrane packing–sensitive dye Flipper-TR. Wild-type α0 decreases membrane order (middle), whereas the F2A/F5A mutant increases it relative to the bare bilayer (right). Scale bar, 5 µm. (**D**) quantification of Flipper-TR lifetimes (ns) for each condition Two-tailed Student’s t-test with Welch’s correction for unequal variances. (**E**) The left micrograph shows membrane constriction and fission of the NT (in magenta) by the Hofund α0 peptide (10 µM). Scale bar, 4 µm. Line-scan analysis (right) revealing progressive radius reduction until fission occurs.

Fluorescence lifetime measurements using the packing-sensitive dye Flipper-TR showed that Hofund increases lipid disorder via a bilayer-couple mechanism (*46*), consistent with leaflet-asymmetric insertion (Fig. 3C,D). By contrast, F2A/F5A increased membrane order, indicative of superficial binding that restricts lipid mobility. By coupling membrane insertion with lipid disorder and curvature generation, these interactions may lower the energetic barrier for fission. Notably, purified Hofund was sufficient to constrict NTs and trigger fission (Fig. 3E), whereas the mutant F2A/F5A helix was not (fig. S3). These results identify Hofund as the molecular trigger of membrane fission through deep asymmetric insertion and raise the question of whether this mechanism is conserved in eukaryotes.

### Membrane fission is performed in eukaryotic ESCRT-III subunits through Hofund elements

We next investigated whether eukaryotic ESCRT-III paralogs might also retain a functional analog of Hofund capable of driving membrane destabilization and fission. We searched for amphipathic helices containing aromatic residues facing the same surface as Hofund elements in the eukaryotic A-type subunits evolutionarily related to Heimdall ESCRT-IIIA: Vps2, Vps24, and Did2 (Fig. 4A). Note that we excluded the non-canonical ESCRT-III subunit, Ist1, from this analysis because in published structures it does not interact directly with the membrane (*34*). Sequence alignments across eukaryotes revealed conserved N-terminal aromatic residues (phenylalanines and tryptophans) in these subunits (Fig. 4B). Among them, Did2, the last ESCRT-III subunit in the sequence and generates the highest curvature (*12*, *34*), retains the most Hofund-like arrangement with two phenylalanines that were previously shown to insert into the hydrophobic core of the membrane (*45*).

**Figure 4.**
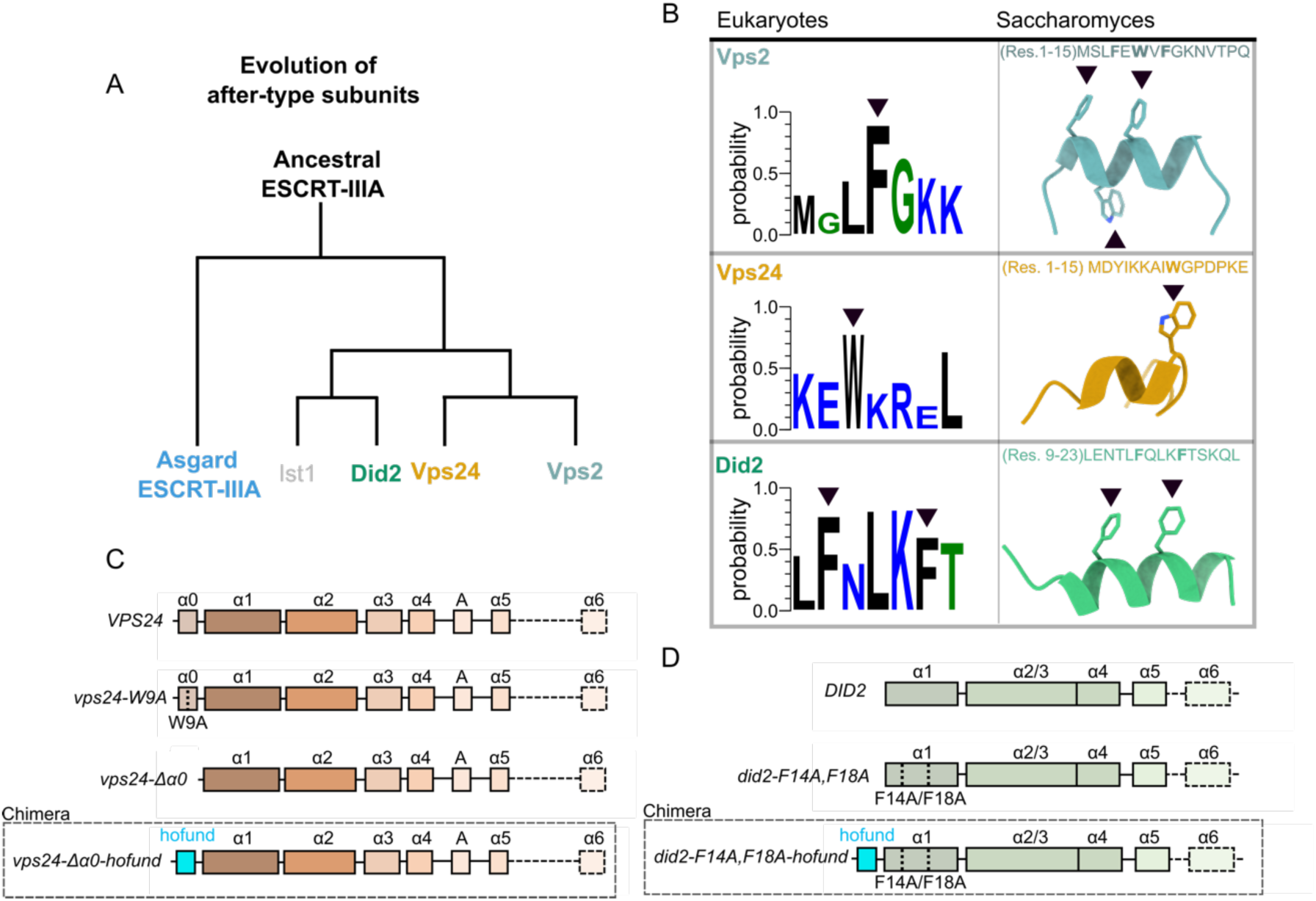
Hofund elements are conserved among eukaryotic A-type subunits. (**A**) Conceptual model, which illustrates the phylogenetic relationship of yeast A-type subunits Did2, Vps24, Did4, and Ist1 and Asgard Heimdall ESCRT-IIIA. (**B**) Amino acid sequence conservation logos (left) and AlphaFold3 structural models (right) of the N-terminal regions of Did2, Vps24, and Did4, highlighting conserved Hofund aromatic residues (black arrowheads). (**C**) α helix domain organization of yeast A-type ESCRT-III subunits Did2 (green) and (**D**) Vps24 (brown), with the mutant constructs designed and used in this study indicated: *did2-F14A/F18A*, *vps24-W9A*, and *vps24-Δα0*. Chimeric variants containing the Heimdall Hofund α0 helix are shown in cyan.

These observations suggest that gene duplications during eukaryogenesis diversified an ancestral ESCRT-IIIA protein, distributing the original Hofund across multiple A-type paralogs. We first tested whether these eukaryotic Hofund elements affect membrane properties *in vitro* using purified N-terminal peptides from the yeast subunits (Fig. 4B). Vps24 is the only A-type subunit in which the Hofund element lies within the α0 helix, as in ESCRT-IIIA, but its α0 folded less efficiently than Heimdall Hofund (fig. S4A) despite its strong hydrophobicity (0.560) and hydrophobic moment (0.418). The Hofund elements in Did4 (Vps2) and Did2, which reside at α1, failed to fold as α helices, suggesting that their structure depends on the full protein core (fig. S4B,C). In the MD simulations, all Vps2, Vps24, and Did2 Hofund elements, even when added pre-folded as amphipathic helices, failed to insert into the bilayer (fig. S4D-F). Consistent with these structural features, none of them induced membrane remodeling or fission *in vitro* (fig. S5).

A defining feature of ESCRT-III subunits is their tendency to form heteropolymers. We hypothesized that a complete Hofund in eukaryotes is reconstituted when A-type subunits copolymerize, rendering each individual element essential for full activity. However, mixing the three peptides together did not trigger fission (fig. S5), likely because oligomerization requires the protein cores (*47*, *48*). To determine how individual Hofund elements contribute to ESCRT-III-mediated fission *in vivo*, we turned to *Saccharomyces cerevisiae*. We generated yeast strains carrying point mutations substituting conserved phenylalanine or tryptophan residues with alanines in Vps24 and Did2 (Fig. 4B-C). Because Vps24 is the only paralog retaining the Hofund element in the α0 helix, we tested its role also by generating a strain lacking the entire helix (Fig. 4C). Vps2 mutants could not be tested due to difficulties encountered when trying to express the protein. The mutant yeast showed a mild defect in the early log phase growth and then stopped growing after the diauxic shift when wild-type cells transition to ethanol metabolism (fig. S6A), which is reminiscent of Snf7 mutants (*49*). Consistently, the mutants failed to grow on ethanol as the carbon source (fig. S6B).

To assess the role of Hofund elements in membrane fission, we followed the trafficking of the amino-acid transporter Mup1 into the vacuole during ILV formation. In wild-type cells, Mup1-GFP is delivered into the vacuole lumen upon methionine addition, whereas Mup1-GFP was blocked at the vacuolar membrane on fission-defective phenotypes (Fig. 5A) (*50*, *51*). All mutants showed Mup1 retention at the endosomal and vacuolar membranes, indicative of a defect in membrane fission (Fig. 5B,C). The *vps24-W9A* showed a partial defect, whereas deleting the entire α0 helix (*vps24-Δα0*) caused a severe block, indicating that the helix itself is essential. Mutating the Hofund residues in Did2 (*did2-F14A/F18A*) produced a strong phenotype comparable to the *vps24-Δα0* helix mutant (Fig. 5B,C). Together, these results demonstrate that eukaryotic Hofund elements are critical for ESCRT-III-mediated membrane fission. These findings support a conserved mechanism in which membrane insertion and destabilization are distributed across subunits and cooperatively reassembled within the ESCRT-III heteropolymer. Furthermore, these results imply that the same mechanochemical principle underlying fission on positively curved NTs *in vitro* applies to the negatively curved necks during ILV formation *in vivo*.

**Figure 5.**
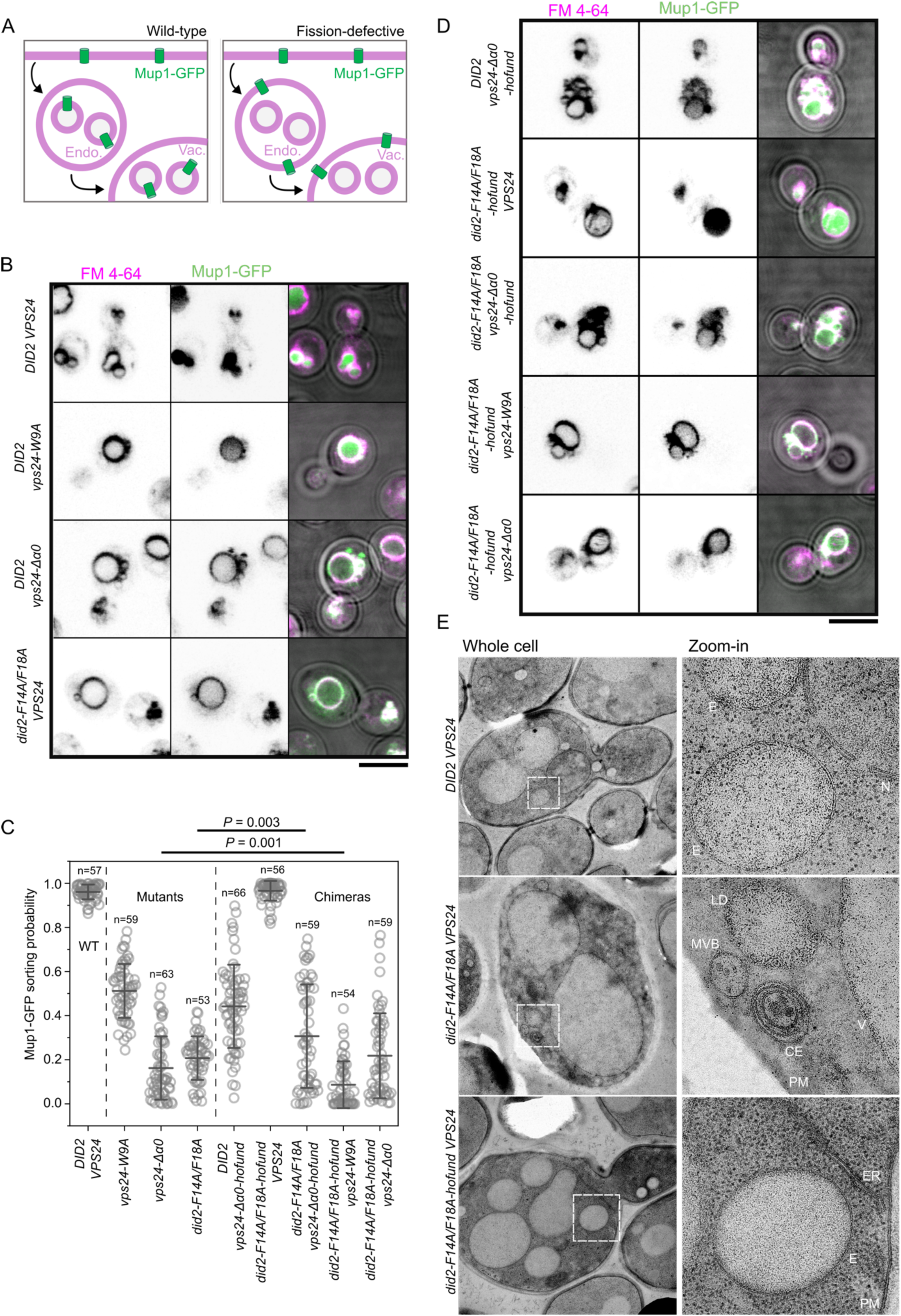
Hofund restores fission efficiency when inserted at the end of the eukaryotic assembly sequence. (**A**) Schematics of Mup1-GFP localization across the endolysosomal pathway in wild-type cells (left) and ESCRT-III-fission defective cells (right). (**B**) Fluorescence micrographs of Mup1-GFP (green) trafficking in wild-type and mutant yeast strains upon methionine addition. FM4-64 (magenta) labels vacuolar membranes. Scale bar is 4 µm. (**C**) Quantification of Mup1-GFP vacuolar lumen as to the Mup1-GFP accumulated at the vacuolar membrane for the indicated strain. N=3 independent experiments. Two-tailed Student’s t-test with Welch’s correction for unequal variances. (**D**) Fluorescence micrographs of Mup1-GFP (green) trafficking in chimeric yeast strains upon methionine addition. FM4-64 (magenta) labels vacuolar membranes. Scale bar is 4 µm. (**E**) EM micrographs of cell cryo-thin sections of wild-type (*DID2 VPS24*), Did2 mutant (*did2-F14A/F18A VPS24*), and Did2 chimera (*did2-F14A/F18A-Hofund VPS24*). E:endosomes; N:nucleus; LD:lipid droplet; MVB:multivesicular body; CE:class E compartment; V:vacuole; PM:plasma membrane; ER:endoplasmic reticulum.

### Hofund at the final step of sequential eukaryotic ESCRT-III assembly is sufficient for ILV formation

If eukaryotic ESCRT-III-mediated membrane fission is triggered by the Hofund elements at the N-termini of the A-type subunits, we reasoned that Heimdall Hofund might act as a mechanical trigger of membrane fission when inserted into the eukaryotic assembly sequence. To test this, we engineered chimeric constructs (*did2-F14A/F18A-hofund* and *vps24–Δα0-hofund*) in which the Heimdall Hofund α0 helix was fused to the eukaryotic Hofund element mutants that are defective in ILV formation (Fig. 4C,D). The *vps24-Δα0-hofund* chimera did not restore ESCRT-III function and resembled the *vps24-W9A* phenotype (Fig. 5C,D). Remarkably, however, introducing Hofund into the Did2 mutant fully restored Mup1 trafficking to wild-type levels, revealing Hofund as the conserved membrane fission trigger and supports that Did2 is the terminal subunit in the eukaryotic assembly sequence (Fig. 5C,D). Thin-section electron microscopy further showed that Hofund fully restored normal endo-lysosomal morphology in *did2-F14A/F18A* cells, eliminating the class E compartments characteristic of impaired fission in the *did2-F14A/F18A* mutant (Fig. 5E) (*5*, *52*).

We also assessed whether a single Hofund could compensate for the combined loss of eukaryotic Hofund elements. The *vps24-Δα0-hofund* chimera provided a modest rescue of the *did2-F14A/F18A* fission defect. However, in the opposite arrangement *did2-F14A/F18A-hofund* failed to rescue the *DID2 vps24-W9A* and *DID2 vps24–Δα0* phenotypes (Fig. 5C,D), indicating that Did2 cannot substitute for defects in the upstream ESCRT-III subunits. While these observations suggest that there is some redundancy in the system, fission strictly requires the correct function of upstream subunits, as previously shown *in vitro* (*12*). In this framework, Did2 likely acts to impose the highest curvature of the filament and surrounding membrane and facilitates the final fission step during ILV formation by the Hofund element. Finally, we asked whether the *did2-F14A/F18A-hofund VPS24* strain, which fully rescued Mup1-GFP trafficking, also restored ethanol-metabolism phenotypes. The chimera failed to recover wild-type growth (fig. S7). Thus, while Hofund-mediated membrane fission has been conserved over 2 billion years (*53*), ESCRT-III subunits present in eukaryotic cells may perform additional more specialized functions.

## Conclusion

Intracellular membrane dynamics, sustained by proteins that remodel membranes through fusion and fission are a hallmark of eukaryotic complexity. Here, we address a long-standing question: does the ancient ESCRT-III machinery, present across all domains of life and likely originating in LUCA, mediate membrane fission outside eukaryotes, or was membrane fission a eukaryotic innovation that emerged with the endomembrane system? Our results provide the first direct evidence that ESCRT-III proteins from Heimdall Asgard archaea, which predate eukaryogenesis, are inherently active in membrane fission. Recent identification of intracellular vesicular structures in Heimdall species (*54*) suggests that ESCRT-III and Hofund may participate in primitive compartmentalization and offers exciting questions about the origins of the endomembrane system.

By using a fully fission-competent Asgard ESCRT-III system, we have resolved a central challenge in the ESCRT-III field. Although major progress has clarified how ESCRT-III filaments assemble, exchange subunits, and generate curvature, the mechanical trigger for fission has remained elusive for more than two decades. Here, we identify an N-terminal amphipathic helix in Heimdall ESCRT-IIIA, Hofund, as the essential fission trigger, consistent with models in which amphipathic helices promote membrane fission (*55*). Hofund binds to lipid bilayers, inserts into the hydrophobic core of the membrane, and destabilizes it through a bilayer-couple mechanism leading to membrane fission, while the rest of the ESCRT-III polymer regulates this activity through curvature generation and ATP-dependent subunit turnover by *ah*Vps4. In bacteria, certain ESCRT-III homologues such as PspA or Vipp1, which contain aromatic residues at their N-terminus, participate in membrane stress-response pathways or thylakoid biogenesis with no direct evidence of membrane fission (*56*). Critically, in these cases, they operate without a dedicated Vps4-type ATPase. Thus, the ability of ESCRT-III to remodel and sever membranes, central to its function in archaea and eukaryotes, appears to be an evolutionary innovation that arose after bacterial and archaeal lineages diverged, likely concomitant with the emergence of the ATPase-driven filament-turnover mechanism.

By testing Hofund *in vitro* and *in vivo*, we demonstrate that it is both necessary and sufficient for membrane fission across opposite curvature contexts: it drives fission on positively curved NTs *in vitro* and also enables ESCRT-dependent fission inside negatively curved membrane necks *in vivo*. This symmetry suggests that the core mechanochemical principle of fission is curvature-agnostic, and that the long-standing *in vitro* and *in vivo* curvature preferences may reflect filament orientation rather than distinct mechanisms.

We show that eukaryotic ESCRT-III subunits retain Hofund elements, but these have become fragmented across multiple paralogs. Their essential roles in ILV formation demonstrate that an ancestral amphipathic-insertion mechanism has been preserved, even as ESCRT-III diversified during eukaryogenesis. Our data support a scenario in which ESCRT-IIIA duplicated and subfunctionalized into Vps2, Vps24, and Did2, acquiring a sequential assembly program in which Did2 provides the terminal fission activity, while the potential Ist1 Hofund element contribution remains unknown. Over evolutionary time, adaptive and non-selective forces likely shaped this diversification (*57*). A potent, sharp Hofund may have limited the ESCRT-III versatility in the eukaryotic ancestor, while the complexification of ESCRT-III could have provided eukaryotic cells with the ability to regulate assembly, membrane curvature generation, and fission activity, which may have opened multiple evolutionary routes that give eukaryotes their rich cellular compartmentalization.

## Methods

### Gene cloning, protein expression, purification, and chemical labeling

Comparison of the Heimdallarchaeota AB_125 ESCRT-IIIA sequence with homologs from representative Asgard phyla (Heimdallarchaeota, Promethearchaeota, Thorarchaeota, Wukongarchaeota, and Hermodarchaeota) revealed that its original UniProt entry (A0A1Q9PC75) lacked five N-terminal residues.

The coding sequences for the full-length Heimdallarchaeota ESCRT-IIIA and for a variant lacking the first 5 residues (ΔN-ESCRT-IIIA) were synthesized and inserted into pET-28a (Novagen) in-frame with an N-terminal His₆–bdSUMO tag (83,84) (Brachypodium distachyon SUMO) to facilitate purification and tag removal. Plasmids were transformed into E. coli Rosetta(DE3)pLysS (Novagen). Cultures were grown in 2×TY medium at 37 °C to mid-log phase (OD₆₀₀ ≈ 0.6), and protein expression was induced with 0.5 mM IPTG for 4 h at 37 °C. Cells were harvested by centrifugation (5,000 g, 10 min, 4 °C), resuspended in buffer A (20 mM Tris-HCl pH 8.0, 500 mM NaCl, 5% glycerol), and lysed by sonication. Lysates were clarified by centrifugation (30,000 g, 30 min, 4 °C) and applied to a HisTrap HP 1 mL Ni²⁺-affinity column (Cytiva). Bound protein was eluted using a linear imidazole gradient (0–500 mM in buffer A).

The His-bdSUMO tag was removed by overnight digestion with bdSENP1 protease at 4 °C, leaving the native N terminus. The cleaved sample was passed over the same Ni²⁺ column to remove protease, tag, and uncleaved protein. The flow-through containing untagged ESCRT-IIIA was further purified by size-exclusion chromatography (Superdex 200 Increase 16/600, Cytiva) equilibrated in buffer A.

For fluorescence imaging, purified proteins were labeled site-specifically at the N terminus using either TFP–Alexa Fluor 488 or Alexa Fluor 568 NHS ester (Thermo Fisher) following the manufacturer’s protocol. Excess dye was removed by desalting (PD-10 column, Cytiva).

### Preparation of lipid-coated silica beads

Supported lipid bilayers (SLBs), giant unilamellar vesicles (GUVs), and membrane nanotubes (NTs) were generated from lipid films deposited onto 40 µm silica beads (Microspheres–Nanospheres), following an adapted version of the bead-hydration method (*58*). Lipid mixtures contained DOPC:DOPS:Atto-647N–DOPE at 59.95:40:0.05, or DOPC:DOPS:DOPE:Atto-647N–DOPE at 20:40:40:0.05 mol%, and were dissolved in chloroform at 1 mg/mL. All lipids were purchased from Avanti Polar Lipids.

Lipid solutions were dried under vacuum for ≥2 h to remove solvent and yield a uniform film. Films were hydrated in 25 mM HEPES-NaOH, pH 7.4, and gently resuspended to form multilamellar vesicles. 10 mL of the vesicle suspension was mixed with 1 mL of silica beads, divided into droplets on parafilm, and dried again under vacuum for ≥1 h to allow complete removal of the aqueous phase. The resulting lipid-coated beads were stored dry and hydrated immediately before use to generate GUVs or to pull lipid nanotubes.

### GUV and NT experiments

GUVs were generated from lipid-coated silica beads as described above. Briefly, lipid-covered beads were hydrated in 1 M trehalose for 15 min at 60 °C in a custom humidity chamber, then transferred to the imaging chamber. At this stage, the newly formed GUVs remained attached to the silica surface; to obtain freely suspended GUVs, the chamber was gently agitated for ∼1 min to promote bead detachment.

Desired purified proteins, specified in each section, were added after GUV release, at the concentrations indicated for each experiment. For protein-encapsulation assays, lipid-coated beads were hydrated directly in buffer containing Heimdall ESCRT-IIIA (1–5 µM) for 30 min at 25 °C before dilution (≥1:50).

For assays in which protein bound from the outside of the tube, NTs were instead generated by gently rolling GUV-coated beads across the chamber surface. In the assays where protein was encapsulated inside the GUVs, NTs were pulled from ESCRT-IIIA-containing GUVs using glass micropipettes fabricated with a P-1000 puller (Sutter Instruments). NTs were formed by contacting the GUV membrane with the micropipette tip and retracting it under micromanipulator control (MP-285, Sutter).

To prevent unwanted tube adhesion during protein-dynamics experiments, glass coverslips were passivated with 2 g/L BSA for 30 min. For NT fission assays, passivation time was reduced to 5 min to allow controlled tube local attachment. Fluorescently labelled protein was always added after NT formation, unless otherwise specified.

All imaging was performed on an inverted spinning-disk confocal microscope (3i Intelligent Imaging Innovation) built around a Nikon Eclipse C1 base, equipped with a 100× 1.49 NA oil-immersion objective and an EM-CCD camera (Evolve, Roper Scientific).

### NT radius calculation

Nanotube radii were quantified using a fluorescence-intensity calibration method (*59*). A flat lipid film of identical composition was deposited on a glass coverslip, and its uniform fluorescence was used to determine the membrane fluorophore density (ρ₀). The radius (R) of each nanotube was then calculated from the total fluorescence per unit length of the tube (Fₗ).

### Quantification of ESCRT-IIIA sorting on membrane NTs

The sorting ratio was defined as the relative membrane area occupied by a single ESCRT-IIIA molecule. For curvature-dependent measurements, the fluorescence intensity of labeled ESCRT-IIIA on nanotubes was normalized to the intensity of the membrane dye (Atto-647N–DOPE) on the same region of interest, and the resulting ratio was plotted as a function of nanotube curvature.

Fluorescence values were obtained from the integrated intensity of line-scan profiles acquired along each nanotube for the protein and membrane channels, respectively, following established procedures. Polarization effects were not corrected, as previously described (*60*).

### ESCRT-IIIA fluorescence intensity quantification

Background fluorescence was quantified as the mean pixel intensity measured from four randomly selected regions of interest (ROIs) lacking ESCRT-IIIA signal, using ImageJ (*61*). The fluorescence associated with individual ESCRT-IIIA puncta was calculated by integrating the signal within a 4 × 4-pixel ROI centered on the fluorescence maximum, after subtraction of the background value multiplied by the corresponding 16 pixels.

### Sample preparation for AFM imaging

GUVs composed of *Escherichia coli* total lipid extract (Avanti Polar Lipids) were generated by electroformation as previously described (*62*). Briefly, 1 mg/mL lipid solution in chloroform was deposited onto indium–tin-oxide–coated (ITO) glass slides and dried under vacuum for ≥1 h. The lipid-coated slides were assembled into a chamber containing 300 mM sucrose and subjected to an alternating electric field (1 V, 10 Hz, 2.7 Vpp at 30 kHz) for 1.5 h to promote GUV formation.

Glass coverslips were plasma-cleaned for 10 min (Diener Electronic GmbH+Co. KG) and immediately overlaid with freshly prepared GUVs mixed with extension buffer (220 mM NaCl, 10 mM HEPES, 2 mM MgCl₂, 10 mM CaCl₂, pH 7.1–7.3) to promote spreading and formation of supported lipid bilayers (SLBs). After 10–15 min incubation, the surfaces were gently rinsed ten times with imaging buffer (25 mM HEPES, 10 mM MgCl₂, pH 6.3) to remove loosely attached vesicles. ESCRT-IIIA and ΔN-ESCRT-IIIA were added to *E. coli* SLBs at final concentrations of 2 µM, in their corresponding buffers immediately prior to imaging.

### Ultrafast atomic force microscopy measurements

Ultrafast AFM imaging was performed on a JPK NanoWizard UltraSpeed system (Bruker) equipped with USC-F0.3-k-0.3 cantilevers (NanoAndMore) with a nominal spring constant of 0.3 N·m⁻¹ and a tip radius <10 nm. Images were acquired in intermittent-contact (AC) mode at a free oscillation frequency of ∼300 kHz and a drive amplitude of 15–20 nm. The sharp cantilever tips ensured high lateral resolution while minimizing tip–sample convolution.

Prior to data acquisition, the deflection sensitivity was calibrated by recording ≥3 force–distance curves on a bare glass substrate, and the spring constant was determined using the thermal noise method (*63*). Scan size, pixel resolution, scan rate, setpoint, and gain parameters were optimized for each sample to minimize tip-induced deformation and ensure stable feedback.

Raw AFM images were processed using JPK SPM Data Processing (v8.4), Gwyddion (v2.65) (*64*), and FIJI/ImageJ (v1.54) (*61*). Filament and ring heights were quantified relative to the underlying lipid bilayer. Briefly, height profiles were extracted from line scans drawn perpendicular to the structures of interest, and the apparent height was calculated as the difference between the peak height of the filament or ring and the mean height of the adjacent flat SLB region. All measurements were performed on flattened (first-order plane-corrected) images to remove background tilt and scanner drift.

### Circular dichroism (CD) spectroscopy

Small unilamellar vesicles (SUVs) composed of 60 mol% DOPC and 40 mol% DOPS were prepared by mixing lipids in chloroform at the desired ratios, evaporating the solvent under argon, and drying the lipid film under vacuum. The dried film was hydrated in 5 mM Na-phosphate buffer (pH 6) and sonicated in a water bath to obtain SUVs. Synthetic peptides corresponding to the Hofund amphipathic helix (MFRDFMGWLLGTKKS), its mutant variant (MARDAMGWLLGTKKS), and the N-terminal helices of Vps24 (MDYIKKAIWGPDPKE), Did4 (MSLFEWVFGKNVTPQ), and Did2 (LENTLFQLKFTSKQL) were obtained from GenScript and dissolved in 5 mM Na-phosphate buffer (pH 6). CD spectra were recorded using 50 µM peptide in the absence or presence of SUVs (250 µg ml⁻¹ final lipid).

Spectra were acquired on a Jasco J-815 circular dichroism spectrophotometer using a 1-mm path-length quartz cuvette. Wavelength scans were collected from 190 to 250 nm with 1-nm steps and a scan speed of 10 nm min⁻¹. Molar ellipticity was calculated with Spectra Manager Analysis software (v2.14.02, Jasco), and α-helical content was estimated following published procedures (*65*). As previously reported for vesicle-containing samples, increased light scattering was observed at wavelengths below ∼200 nm.

### Molecular dynamics simulations

Peptide structures were predicted using AlphaFold3 (*44*) (web server implementation), and all simulations were performed in GROMACS (*66*) v2023.3 using the CHARMM36m force field (*67*).

A lipid bilayer composed of DOPC:DOPS (70:30 mol%) with lateral dimensions of 12 × 12 nm was generated in CHARMM-GUI (*68*). The membrane was equilibrated following the standard six-step CHARMM-GUI protocol, in which positional restraints on lipid headgroups are gradually released while increasing the time step. After equilibration, solvent and ions were removed to obtain the relaxed bilayer, which was used as the starting point for all peptide–membrane systems.

Each peptide was positioned 1 nm above the membrane surface along the bilayer normal (z-axis), and the system was re-solvated with water and ions to neutralize the overall charge and reach 150 mM NaCl. Energy minimization was performed using the steepest-descent algorithm (1000 steps) with positional restraints of 1000 kJ·mol⁻¹·nm⁻² applied to the peptide. A three-stage equilibration followed, progressively releasing peptide restraints and increasing the integration time step. Temperature (310 K) and pressure (1 bar) were maintained with the v-rescale thermostat (*69*) and C-rescale barostat (*70*), respectively.

For each peptide, four independent 500-ns production runs were carried out. Trajectories were corrected for periodic boundary conditions prior to analysis, and the first 300 ns of each trajectory were discarded as equilibration. Peptide insertion depth was quantified using MDAnalysis (*71*) by calculating the z-distance of each residue relative to the phosphate plane of the bilayer; residues located below this surface were assigned negative insertion values. Depth values were averaged across all replicas.

### Confocal FLIM live imaging

Fluorescence lifetime imaging microscopy (FLIM) was performed on a Nikon Eclipse Ti2 inverted microscope equipped with a motorized stage, a point-scanning A1 confocal system, and a 100× 1.45 NA oil-immersion objective (Nikon, MRD01905). Excitation was provided by a pulsed 485-nm diode laser (PicoQuant LDH-D-C-485) controlled by a Sepia PDL-828 driver and operated at 20 MHz.

Images were acquired at 128 × 128 pixels (0.17 µm per pixel) with a pixel dwell time of 2.7 µs, corresponding to a scan speed of ∼100 Hz and a frame time of 126.7 ms. The confocal pinhole was set to 1.4 Airy units. Photons were collected continuously for 60 s per measurement. Emission was passed through a 600/50-nm bandpass filter and detected using a PMA Hybrid 40 detector (PicoQuant). Time-correlated single photon counting (TCSPC) data were recorded using a MultiHarp 150 module. Instrument control and acquisition were carried out in NIS-Elements AR 3.30.05.

### Preparation of GUVs stained with Flipper-TR

GUVs for FLIM experiments were prepared from DOPC and DOPS (Avanti #850375 and #840035) mixed at the indicated molar ratios. Lipids were stored in chloroform (1–10 mg ml⁻¹) under argon at −80 °C. For each preparation, 0.25 mg total lipid was dispensed into glass vials, the solvent was evaporated under a gentle argon stream, and residual chloroform was removed by overnight desiccation in a vacuum oven.

Lipid films were rehydrated in 10 mM HEPES buffer (pH 7.4) to 0.5 mg ml⁻¹ and briefly mixed. Fifty-micrometer silica beads (Sigma #904384-2G) were washed three times in distilled water, diluted 1:3, and deposited onto 2–3 µl lipid droplets placed on parafilm. The lipid–bead mixture was dried again overnight under vacuum to yield lipid-coated beads.

Microscopy chambers were prepared by plasma-cleaning glass coverslips (Harrick #PDC-32G, 2 min, high power) and mounting them onto Ibidi sticky-slides (#80608). Coverslips were passivated with 0.5 mg ml⁻¹ PLL-g-PEG [PLL(20)-g[3.5]-PEG(2); SuSoS] in 10 mM HEPES (pH 7.4) for 30 min at room temperature, followed by three washes in 25 mM HEPES, 10 mM MgCl₂ (pH 7.2).

Lipid-coated beads were gently scraped from the parafilm and rolled across the chamber surface to release GUVs. After 30 min, the chamber was filled with imaging buffer containing 1 µM Flipper-TR (Spirochrome #SC020) and incubated until probe equilibration.

### Flipper-TR lifetime estimation

Flipper-TR is a mechano-responsive molecular probe that changes its fluorescence properties in response to changes in membrane packing. Due to the photochemistry of the molecule, Flipper-TR photon arrival time distribution follows a bi-exponential decay, in the form:, where and represent the amplitude and the decay of a given exponential, respectively. We analzyed lifetime decays using SymPhoTime 64 v2.7 (PicoQuant, Germany). We manually selected GUVs in the field of view using the paint tool to exclude background, and performed 2-exponent reconvolution fits on the ROI selections to obtain the average lifetime (intensity-weighted average lifetime τ_av int_) (*72*). No IRF measurement was used as input, but software generated. No exclusion criteria was applied on GUV based on lifetime values or goodness of fit.

### Yeast growth and strains

*S. cerevisiae* strains were generated in the MKY100 (Kaksonen group). Cells were grown in rich medium (YPD - 1% Yeast extract, 2% Bacto peptone, and 2% Glucose or 4% Ethanol) or synthetic medium (SD - 0.67% Yeast nitrogen base w/o Folic acid w/o Riboflavin, 2% Glucose. 0.192% SC-Trp) and supplemented with appropriate antibiotics. All strains were cultured at 30°C. Gene construction and transformation protocols were performed as established (*73*). To generate mutants, mutant plasmids were generated and integrated into the endogenous locus of *vps24Δ* or *did2Δ* strains. All constructs and strains were verified by PCR and sequencing. All strains used in this study are listed table S1.

**Supplemental table 1:**
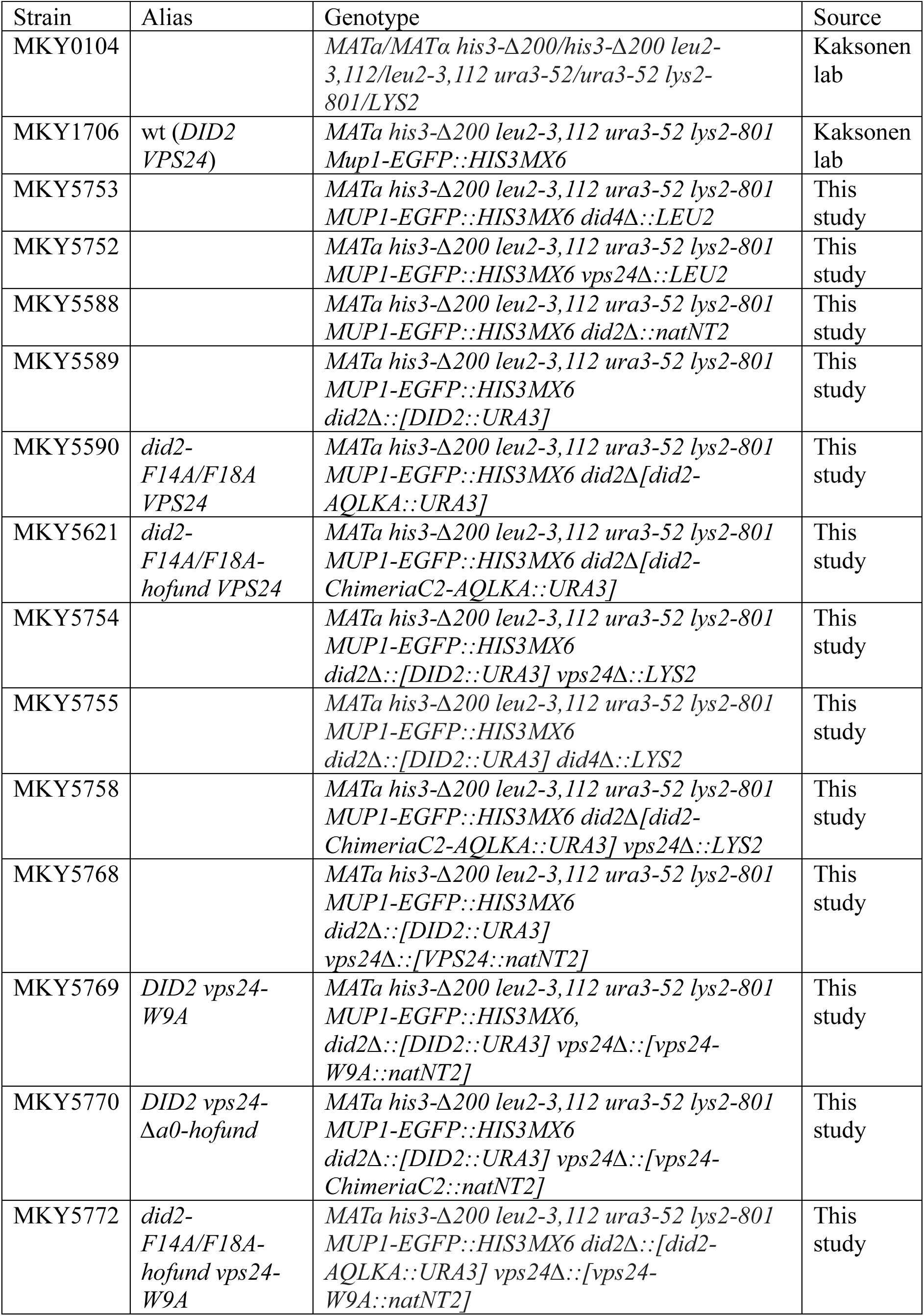
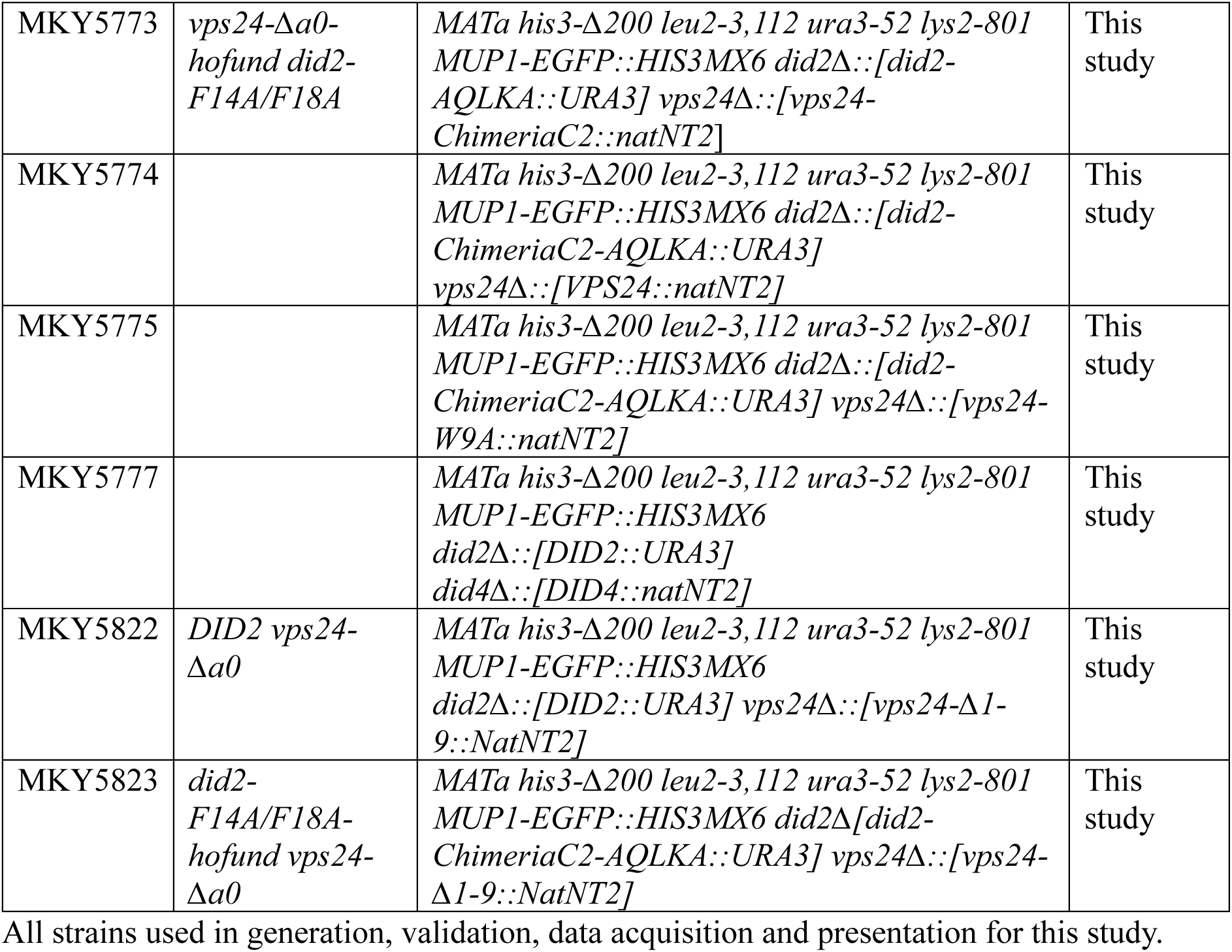
*S. cerevisiae* strains used in this study.

### Mup1-GFP imaging and scoring

Mup1-GFP strains were grown to early log phase in SD -Trp. FM4-64 dye (Invitrogen) was added at the concentration specified by the manufacturer and allowed to internalize for 40 minutes at 30°C and then placed on ice. Cells were image on a Nikon Eclipse Ti2 inverted microscope equipped with a motorized stage, a point-scanning A1 confocal system, and a 100× 1.45 NA oil-immersion objective (Nikon, MRD01905) at room temperature.

Vacuolar localization of Mup1-GFP was quantified as the Mup1-GFP vacuolar sorting probability. Line-scan profiles were generated in FIJI by drawing a 1-pixel-wide line across the widest diameter of FM4-64-labeled vacuoles. FM4-64 intensity peaks defined the vacuolar membrane, and the region between these peaks was assigned as the lumen. For each vacuole, the integrated Mup1-GFP fluorescence intensity was calculated for the lumen (area under the curve between FM4-64 peaks) and for the membrane (sum of intensities at the FM4-64 peaks).

The Mup1-GFP luminal values were normalized to the total GFP signal within the vacuole (summatory of lumen and membrane Mup1-GFP signals).

### Thin section transmission electron microscopy

Yeast cultures were grown in YPD at 30 °C to mid-log phase (OD₆₀₀ ≈ 0.5–0.8) and harvested by gentle centrifugation (500 × g, 3 min). Cell pellets were transferred into 100-µm-deep aluminum carriers (Leica Microsystems) and vitrified by high-pressure freezing using a Leica EM ICE system, following established procedures for preserving endomembrane ultrastructure (*74*).

Vitrified samples were transferred under liquid nitrogen into freeze-substitution medium consisting of 0.1% uranyl acetate in anhydrous acetone and maintained at −90 °C for 42 h in a Leica EM AFS2 freeze-substitution unit. Temperature was then increased stepwise to −25 °C over 12 h. After three washes in anhydrous acetone, samples were infiltrated with increasing concentrations of Lowicryl HM20 (25%, 50%, 75%, and 100%, 2 h each), followed by overnight infiltration in 100% HM20. Polymerization was carried out under UV illumination at −25 °C for 24 h, after which the temperature was gradually raised to 20 °C over 3–4 days to complete embedding (*75*, *76*).

Polymerized blocks were trimmed with a platinum-coated razor blade (Electron Microscopy Sciences) on a Leica UC7 ultramicrotome. A ∼1 mm × 1 mm block face was produced using a Diatome 45° trimming tool and subsequently polished. Ultrathin sections (70 nm) were cut using a Diatome Ultra 35° diamond knife and collected onto 100-mesh carbon-coated formvar grids (Electron Microscopy Sciences).

Grids were post-stained with lead citrate for 30 s (*77*) and rinsed thoroughly with freshly boiled, CO₂-free distilled water. Images were acquired on a Thermo Fisher Talos L120C transmission electron microscope operating at 120 kV, and data were recorded using the integrated CETA 16M camera.

## Supplementary Figures

**Figure S1.**
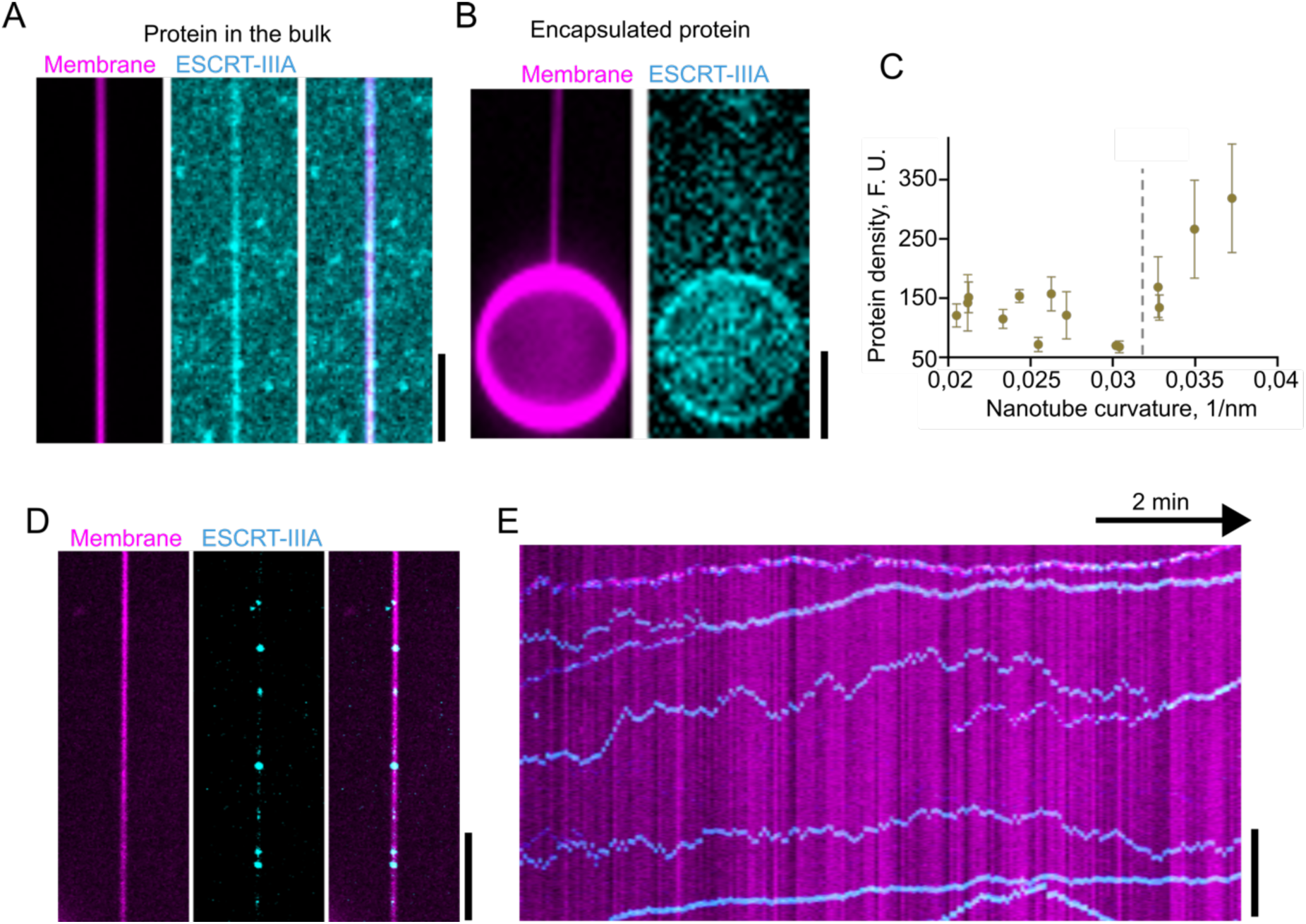
ESCRT-IIIA binds and self-assembles preferentially on positively over negatively curved membranes. (**A**) Initial recruitment of ESCRT-IIIA (cyan) onto a positively curved lipid NT (magenta), visible as a uniform increase in ESCRT-IIIA fluorescence intensity along the NT. Scale bar, 2 µm. (**B**) GUV-encapsulated ESCRT-IIIA fails to bind or self-assemble on the negatively curved interior of a pulled NT. ESCRT-IIIA shows only weak binding to the flat inner GUV membrane, with most protein remaining in the lumen. Scale bar, 3 µm. (C) Curvature-dependent enrichment of ESCRT-IIIA on NTs, quantified as integrated ESCRT-IIIA fluorescence intensity normalized to membrane fluorescence as a function of NT curvature. (D) Formation of discrete ESCRT-IIIA foci (cyan) on the surface of a positively curved NT (magenta). Scale bar, 2 µm. (E) Kymograph showing the dynamic behavior of ESCRT-IIIA foci over time, corresponding to the NT shown in panel D. Scale bar, 2 µm.

**Figure S2.**
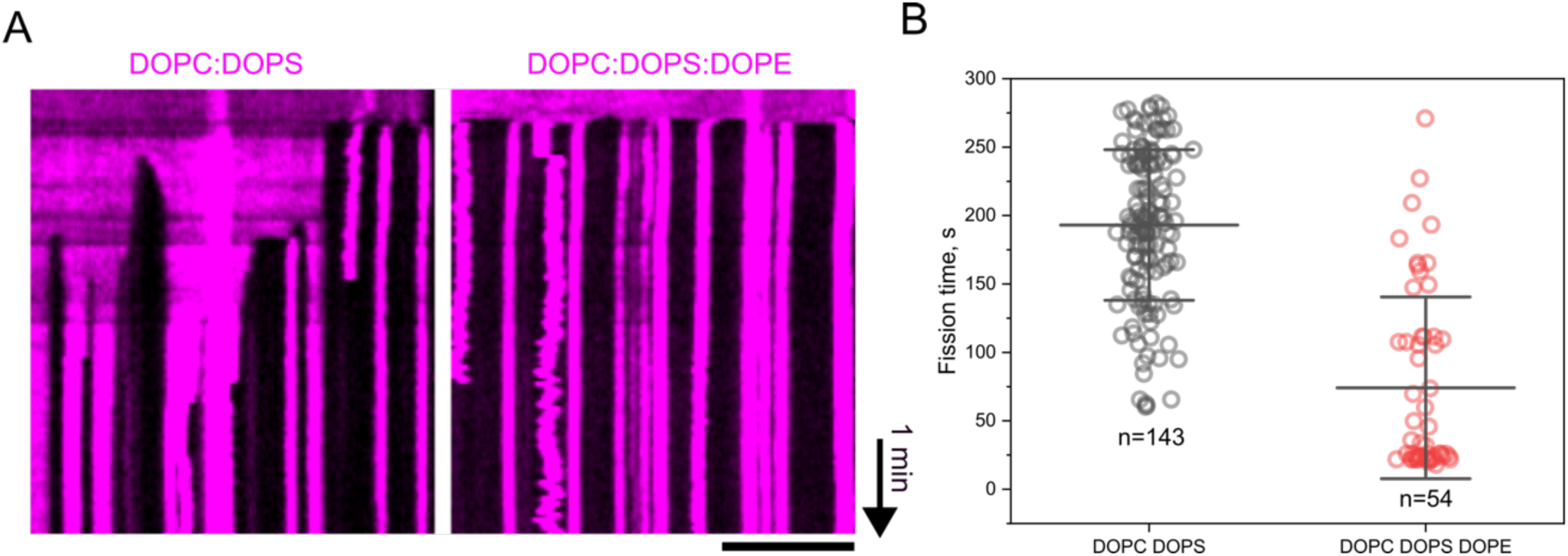
Membrane lipid composition modulates the efficiency of ESCRT-IIIA–mediated fission. (**A**) Kymographs of lipid NTs composed of DOPC:DOPS (60:40 mol%) or DOPC:DOPS:DOPE (20:40:40 mol%) pre-coated with ESCRT-IIIA, followed by addition of free ESCRT-IIIA, *ah*Vps4, and ATP (1 µM / 5 µM / 10 mM, respectively). Enhanced fission frequency is observed in the presence of DOPE. Scale bar, 3 µm. (**B**) Quantification of fission times for the conditions shown in panel A. Two-tailed Student’s t-test with Welch’s correction for unequal variances.

**Figure S3.**
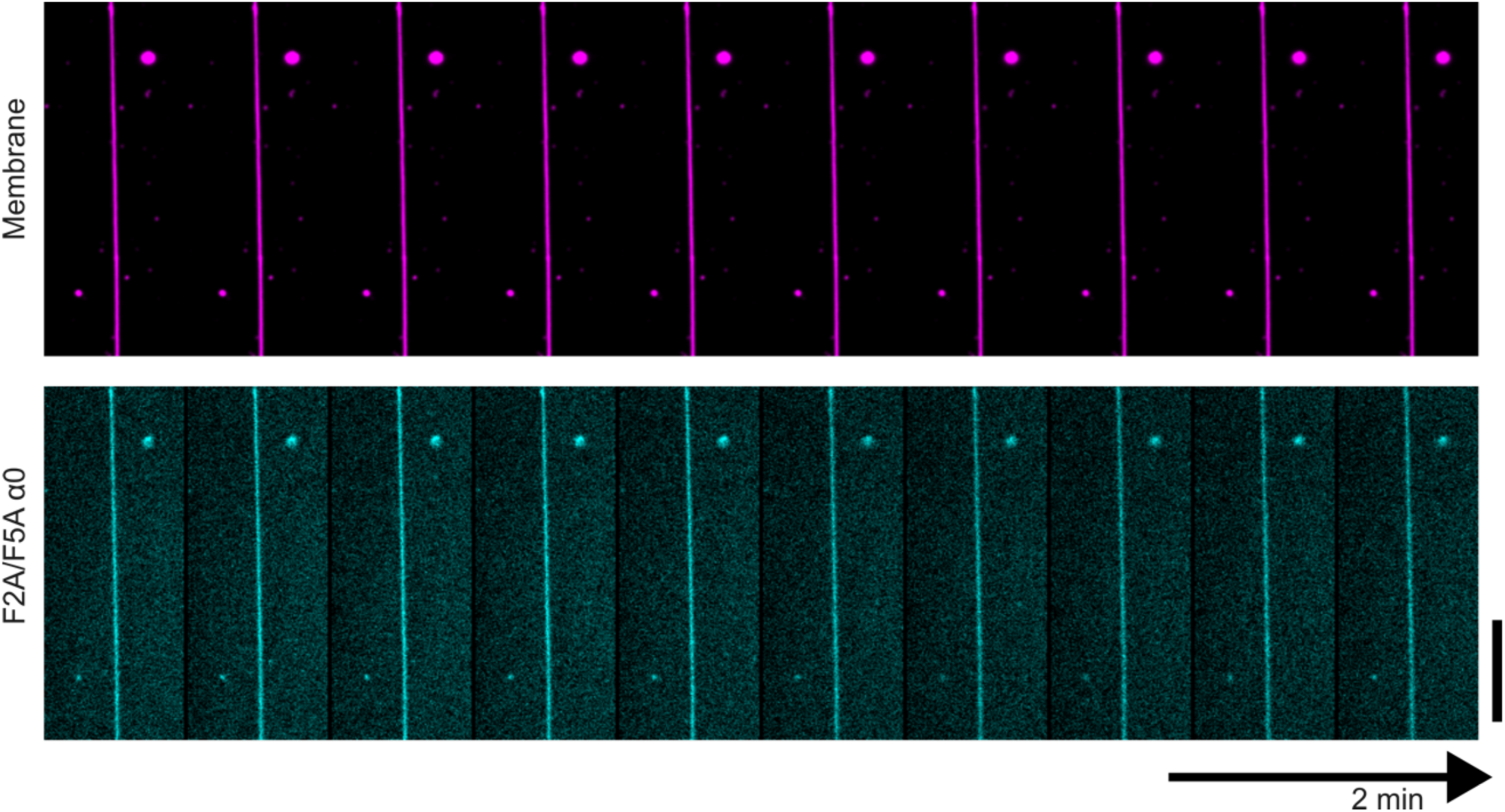
Mutating the phenylalanines in Hofund abolishes its ability to trigger membrane fission. Time-lapse fluorescence micrographs showing that the Hofund F2A/F5A mutant fails to constrict or sever lipid NTs when added at 10 µM, in contrast to the robust fission activity of the wild-type Hofund. Scale bar, 2 µm.

**Figure S4.**
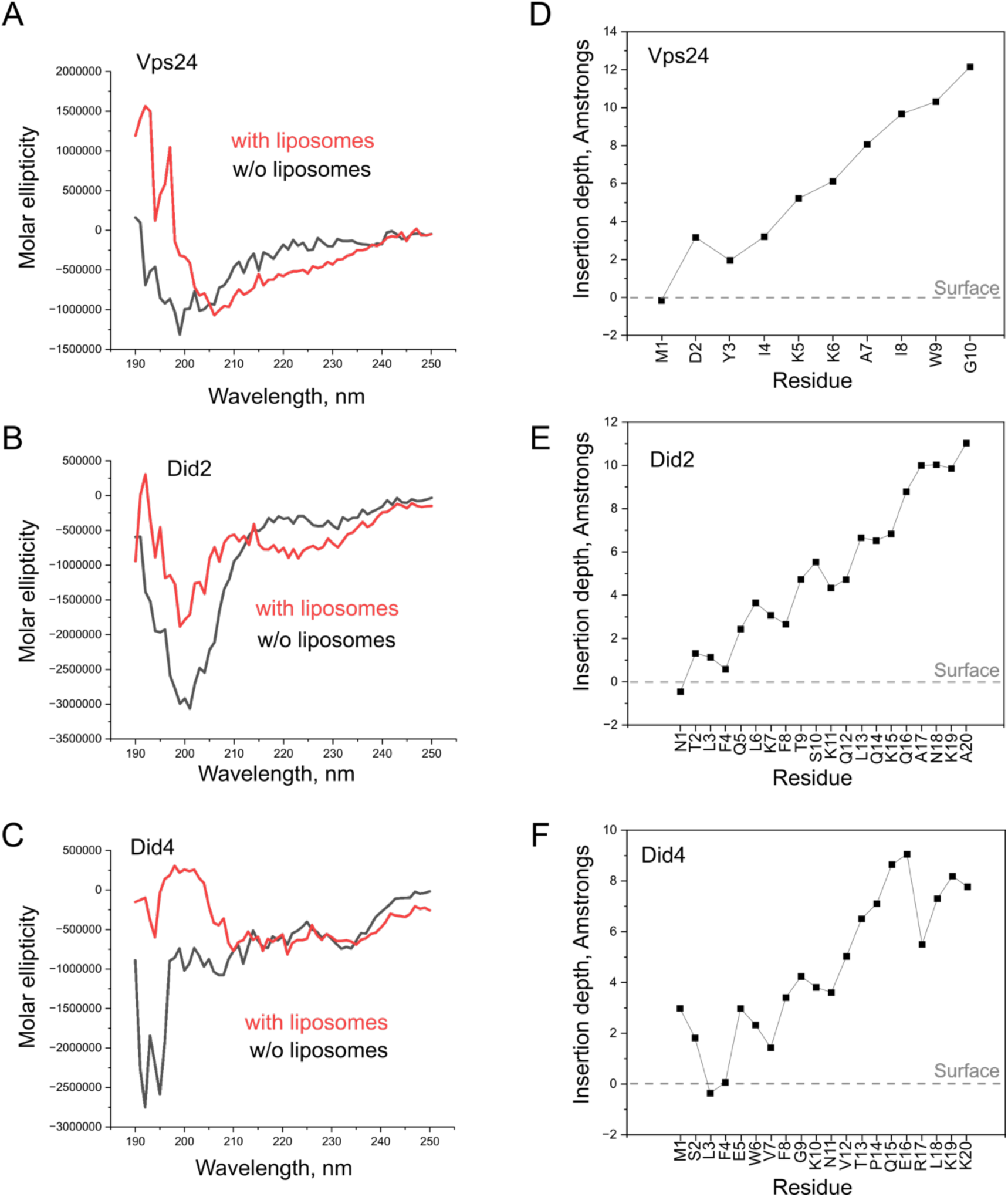
Eukaryotic ESCRT-III N-terminal regions do not individually retain Hofund-like membrane-insertion properties. (**A-C**) CD spectra of the N-terminal regions of Vps24, Did2, and Did4 in the absence (black) or presence (red) of DOPC:DOPS 60:40 mol% membrane liposomes, showing limited or no helix formation upon membrane binding. (**D-F**) All-atom MD simulations of the same N-terminal regions on a DOPC:DOPS 60:40 mol% bilayer reveal negligible membrane insertion compared to Hofund. The gray dashed line indicates the average position of the lipid phosphate plane.

**Figure S5.**
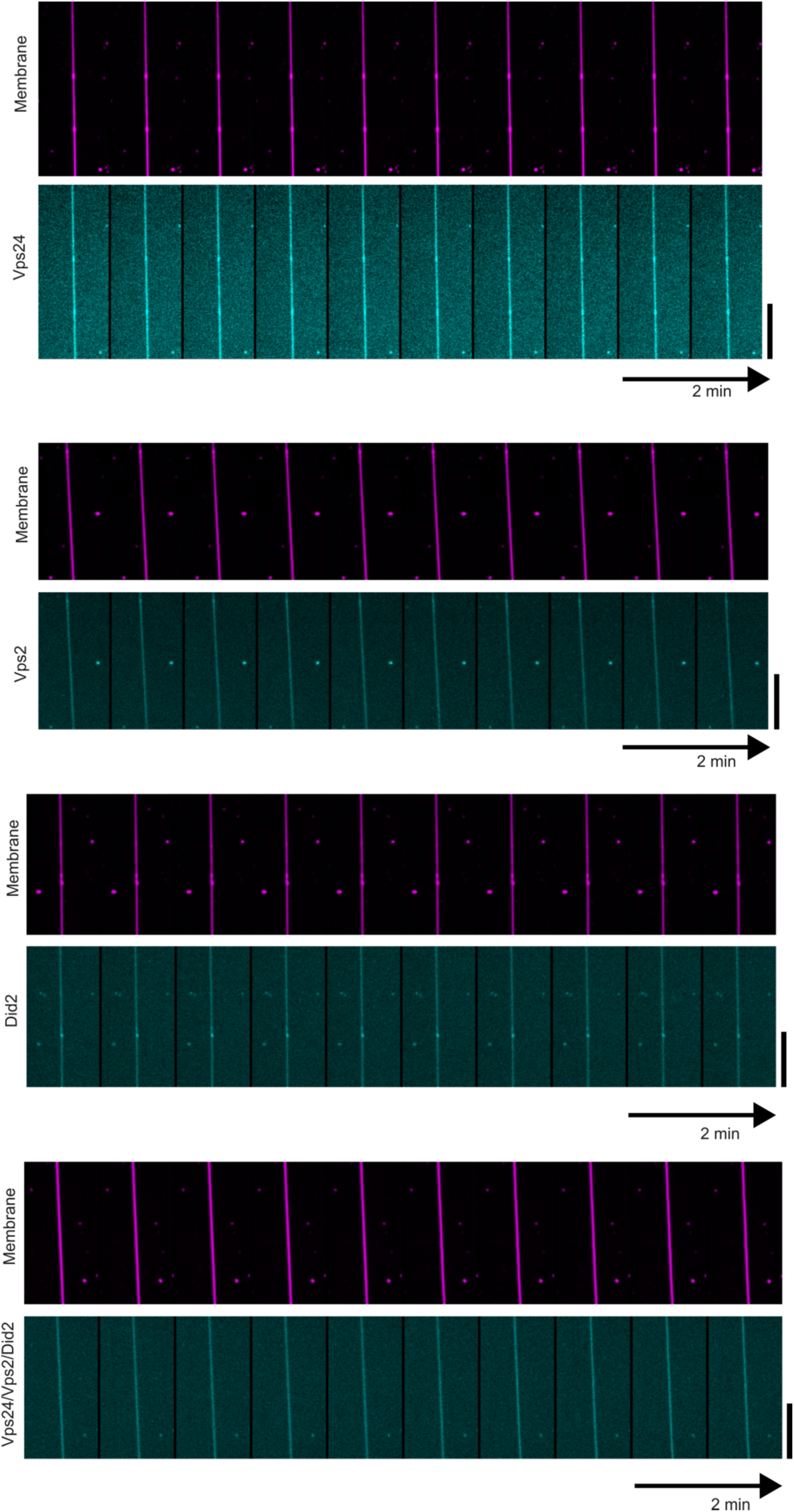
Individual or combined N-terminal peptides of eukaryotic A-type ESCRT-III subunits fail to promote membrane fission. Time-lapse fluorescence micrographs of pre-curved lipid NTs (magenta) after addition of the indicated N-terminal peptides (cyan) at 10 µM. Neither individual peptides nor their combinations trigger membrane constriction or fission, in contrast to the Heimdall Hofund helix. Scale bars, 2 µm.

**Figure S6.**
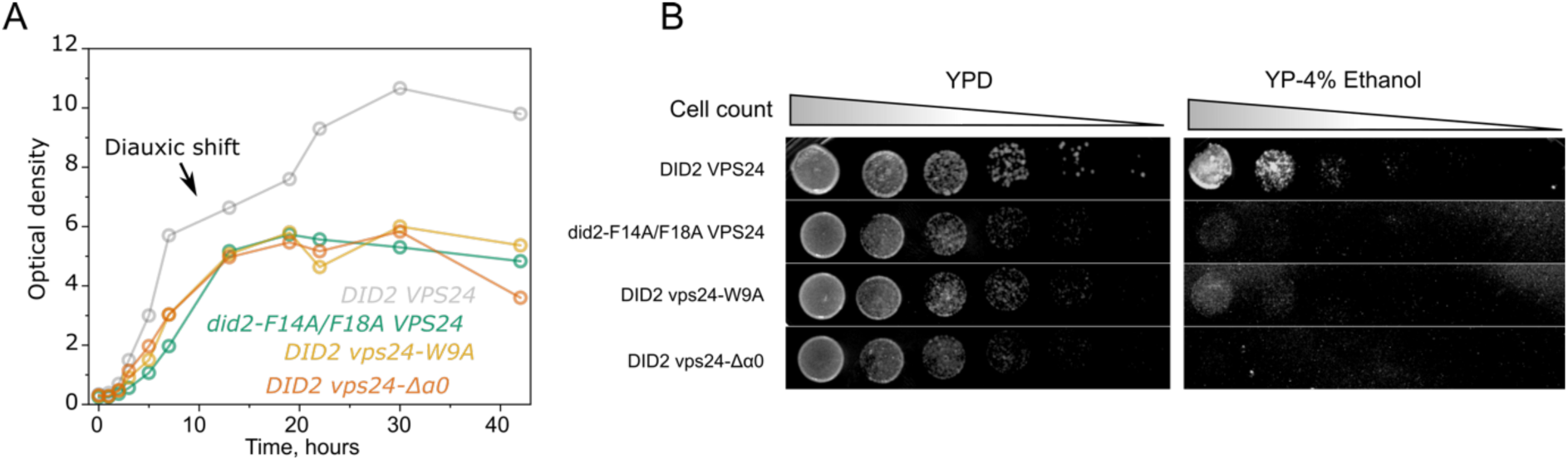
Mutations in eukaryotic Hofund elements impair growth after the diauxic shift. (**A**) Growth curves of wild-type and mutant yeast strains reveal that mutations disrupting Hofund-like aromatic residues cause a failure to resume growth after the diauxic shift, indicating an inability to metabolize ethanol once glucose is depleted. (**B**) Spot assays on YPD and ethanol-based media further confirm that Hofund-mutant strains are unable to grow when ethanol is the sole carbon source.

**Figure S7.**
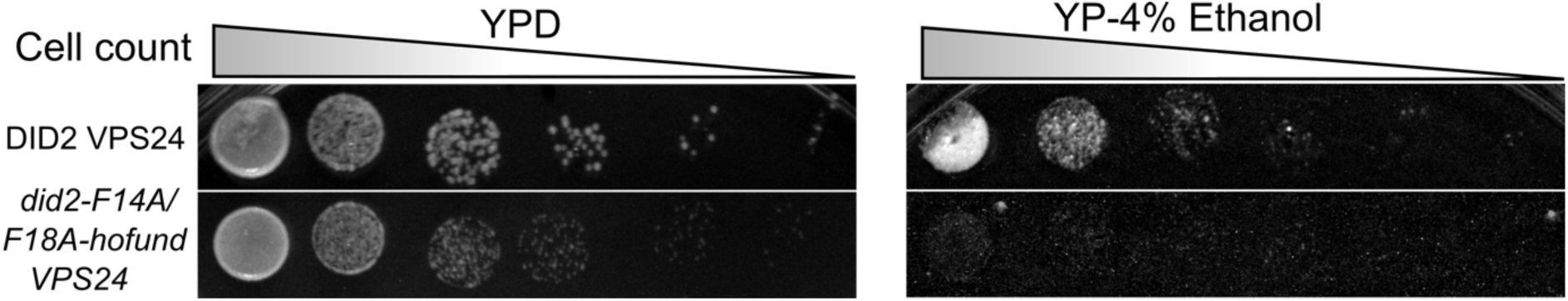
Hofund rescues ILV membrane fission but does not restore metabolic defects. Spot assays on YPD and ethanol-based media show that while introducing Hofund into the Did2 mutant restores ESCRT-III–dependent membrane fission, does not rescue the growth defects associated with ethanol utilization. Mutant strains carrying Hofund-fusion constructs remain unable to grow when ethanol is the sole carbon source, indicating that metabolic defects are independent of membrane-fission activity.

